# Integration of CTCF Loops, Methylome, and Transcriptome in Differentiating LUHMES as a Model for Imprinting Dynamics of the 15q11-q13 Locus in Human Neurons

**DOI:** 10.1101/2024.03.26.586689

**Authors:** Orangel J. Gutierrez Fugón, Osman Sharifi, Nicholas Heath, Daniela C. Soto, J. Antonio Gomez, Dag H Yasui, Aron Judd P. Mendiola, Henriette O’Geen, Ulrika Beitnere, Marketa Tomkova, Viktoria Haghani, Greg Dillon, David J. Segal, Janine M. LaSalle

**Affiliations:** Genome Center, Department of Biochemistry and Molecular Medicine, University of California Davis, Davis, 95616, USA; Department of Medical Microbiology and Immunology, School of Medicine, University of California Davis, Davis, 95616, USA; Department of Psychiatry and Biobehavioral Sciences, University of California Los Angeles, Los Angeles, 90095, USA; Department of Natural Science, Seaver College, Pepperdine University, Malibu, 90263, USA; Max Delbrück Center for Molecular Medicine, Berlin, 13125, Germany; Ludwig Cancer Research Center, University of Oxford, Oxford, OX3 7DQ, United Kingdom; Genetics and Neurodevelopmental Disorders Unit, Biogen, Cambridge, 02142, USA

## Abstract

Human cell line models, including the neuronal precursor line LUHMES, are important for investigating developmental transcriptional dynamics within imprinted regions, particularly the 15q11-q13 Angelman (AS) and Prader-Willi (PWS) syndrome locus. AS results from loss of maternal *UBE3A* in neurons, where the paternal allele is silenced by a convergent antisense transcript *UBE3A-ATS*, a lncRNA that normally terminates at *PWAR1* in non-neurons. qRT-PCR analysis confirmed the exclusive and progressive increase in *UBE3A-ATS* in differentiating LUHMES neurons, validating their use for studying *UBE3A* silencing. Genome-wide transcriptome analyses revealed changes to 11,834 genes during neuronal differentiation, including the upregulation of most genes within the 15q11-q13 locus. To identify dynamic changes in chromatin loops linked to transcriptional activity, we performed a HiChIP validated by 4C, which identified two neuron-specific CTCF loops between *MAGEL2-SNRPN* and *PWAR1-UBE3A*. To determine if allele-specific differentially methylated regions (DMR) may be associated with CTCF loop anchors, whole genome long-read nanopore sequencing was performed. We identified a paternally hypomethylated DMR near the *SNRPN* upstream loop anchor exclusive to neurons and a paternally hypermethylated DMR near the *PWAR1* CTCF anchor exclusive to undifferentiated cells, consistent with increases in neuronal transcription. Additionally, DMRs near CTCF loop anchors were observed in both cell types, indicative of allele-specific differences in chromatin loops regulating imprinted transcription. These results provide an integrated view of the 15q11-q13 epigenetic landscape during LUHMES neuronal differentiation, underscoring the complex interplay of transcription, chromatin looping, and DNA methylation. They also provide insights for future therapeutic approaches for AS and PWS.

## Introduction

Human *in vitro* models play a crucial role in advancing our understanding of neurodevelopmental disorders. These models offer a controlled environment to investigate the intricate interplay of genetic and epigenetic gene regulation, shedding light on the molecular mechanisms underlying these disorders. However, for *in vitro* models of imprinted neurodevelopmental disorders associated with human 15q11.2-q13.3 deletions and duplications, there are additional considerations due to the developmental transcriptional dynamics of this locus in early postnatal neuronal maturation (1). Human *in vitro* models are essential for understanding neurodevelopmental disorders linked to the 15q11.2-13.3 region due to interspecies genetic and epigenetic differences. Specifically, the transcript that silences paternal *UBE3A* in neurons exhibits different splicing and termination points in non-neuronal cells when comparing mice to humans (2). This distinction underscores the necessity of human-specific models to accurately explore the epigenetic landscape and inform therapeutic development.

Angelman syndrome (AS) is a severe neurogenetic disorder affecting approximately 1 in 15,000 births. It is characterized by developmental delay, seizures, language deficiency, ataxic gait, and a happy demeanor (3). AS is caused by a functional loss of *UBE3A* located within the 15q11-q13 region, with most cases arising from a *de novo* maternal allele deletion spanning about 6 million base pairs (4,5). The *UBE3A* gene is subject to biallelic expression in most tissues, meaning that both maternal and paternal alleles are active. However, within neuronal cells, the paternal allele of *UBE3A* is imprinted, which silences its expression and leaves the maternal allele as the sole contributor to the gene’s function in these cells (6). *UBE3A* codes for a ubiquitin ligase E3A protein which is essential for synaptic development (7,8).

*SNRPN* is located in the forward strand opposite strand to *UBE3A* which is transcribed from the reverse strand and encodes a protein regulator of alternative splicing (9,10) (**Figure 1**). The *SNRPN* protein coding region is at the 5’ end of a longer 700 kb transcript that includes an extensively spliced, long non-coding RNA (lncRNA) (11). In neurons and non-neurons, paternal expression of this lncRNA begins at the *SNRPN* promoter and extends past *SNORD116* and *SNORD115*, a repetitive region of small nucleolar RNAs (snoRNA) that are processed from the larger host gene transcript (*SNHG14*). The Prader-Willi syndrome imprinting center (PWS-ICR) regulates gene expression on the paternal chromosome, and the deletion of *SNORD116* within this region is crucially linked to the development of Prader-Willi syndrome (12,13). In non-neurons the transcript terminates at the non-coding *PWAR1*, which is an exon within *SNHG14* (14). However, in neurons, *SNHG14* transcription continues beyond *PWAR1* through the *SNORD115* cluster and further extends antisense to *UBE3A (UBE3A-ATS)*. This antisense transcript has been shown to be responsible for the silencing of the paternal allele in neurons (15).

**Figure 1.**
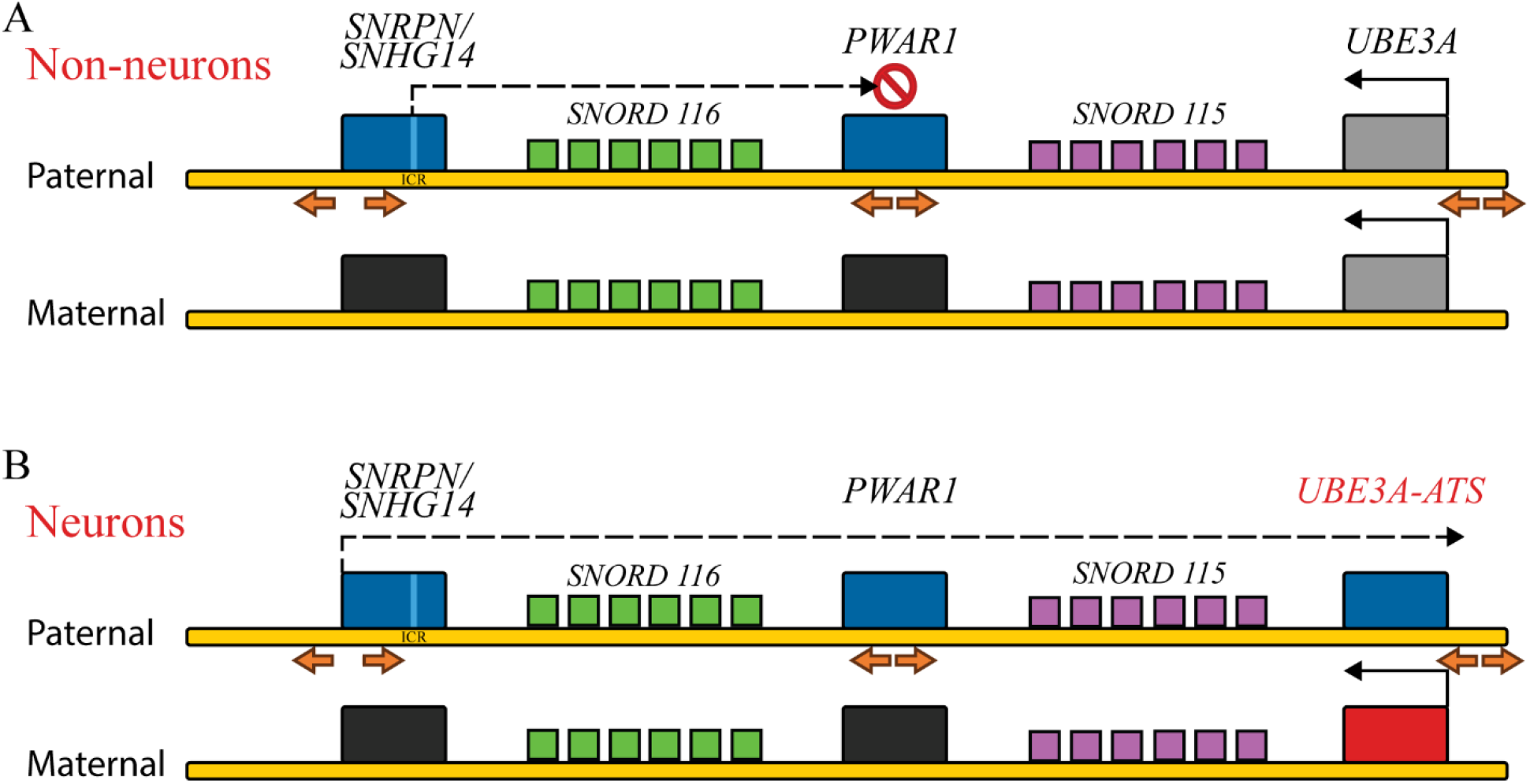
Simplified map of the human 15q11-q13 imprinted locus showing transcription initiation and termination of *SNHG14* lncRNA in: **A.** non-neurons, compared to extension through *UBE3A-ATS* in **B.** neurons. Red indicates exclusively maternally expressed genes, blue is exclusively paternally expressed genes, gray is biallelically expressed and black is repressed. Arrows indicate divergent CTCF binding sites. *SNORD116* repeats are shown in green, while *SNORD115* repeats are shown in purple. Light blue band on *SNRPN* gene represents PWS-ICR.

The sequential molecular events that lead to developmental regulation of transcript elongation remains a central question in the epigenetics of the 15q11-q13 region. Non-neuronal cells from a PWS patient with a *SNORD116* deletion that included *PWAR1* were shown to express *UBE3A-ATS,* suggesting the existence of a boundary region (14). The presence of binding sites for the insulator protein CTCF (CCCTC-Binding Factor) at *PWAR1* has led to the hypothesis that this boundary may serve as the barrier to transcriptional extension in non-neurons (16), but the role of chromatin topology has not been assayed explicitly.

CTCF is a strong candidate for the transcription regulation dynamics in the 15q11-q13 region based on its established function at other loci. CTCF associates with cohesin to form chromatin loops which have been shown to regulate tissue and allele-specific differential gene expression (17,18). Reduced CTCF binding correlates with CpG hypermethylation at its canonical binding motif (19). The CTCF binding motif is recognized by the CTCF protein, which can bind to DNA and influence gene expression in both the upstream and downstream directions from its binding site (20). Chromatin loops are formed preferentially by two convergent CTCFs and a stabilizing cohesin ring (20). Cohesin initially binds and begins to extrude chromatin but tends to stop when it encounters convergent CTCF dimers (20,21). This has been shown to be a cyclical and dynamic process with CTCFs binding and unbinding in a matter of several seconds while cohesin can remain bound to chromatin for several minutes (22). Previous studies have suggested the existence of a neuronal transcriptional collision mechanism in which the *UBE3A-ATS* silences paternal *UBE3A* in neurons by outcompeting the *UBE3A* sense transcript, but the exact mechanism is poorly understood in relation to CTCF and chromatin topology (23,24).

A major challenge to the field is that no *in vitro* model can fully replicate the dynamic processes that occur during neurodevelopment in the human brain. Differentiation protocols might not accurately recapitulate the complex maturation steps that *UBE3A-ATS* expressing neurons undergo *in vivo*. Moreover, epigenetic modifications crucial for the regulation of *UBE3A* expression may not be fully established or maintained in these *in vitro* systems. Models for studying the AS/PWS locus include SH-SY5Y cells and human induced pluripotent stem cells (iPSCs) from AS patients (16). However, SH-SY5Y are aneuploid and derive from cancer cells and thus may have an aberrant epigenetic profile. While patient-derived iPSCs hold great promise, full differentiation to mature neurons is a challenging and inconsistent process that can extend beyond seven weeks (16). Despite their valuable insights, these models might not fully capture the intricate epigenetic complexities inherent in the 15q11.2-q13.3 locus and other disease loci with complex neuronal expression patterns.

In contrast, the human LUHMES (Lund human mesencephalic) cell line may be an ideal model to study neurodevelopmental disorders with an epigenetic component. LUHMES cells are female human embryonic neuronal precursor cells capable of sustained proliferation, which is attributed to the presence of an engineered tetracycline-inducible (Tet-off) v-myc transgene (25). When exposed to tetracycline along with glial cell-derived neurotrophic factor (gDNF) and dibutyryl cAMP, these cells can undergo differentiation into postmitotic dopaminergic neurons displaying the presence of β-tubulin, synaptophysin and the enzyme tyrosine hydroxylase. Furthermore, LUHMES cells display spontaneous electrical activity inherent to neurons (25). Compared to pluripotent stem cell lines, they are relatively easy to grow and differentiate into neurons within one week. However, this short life span can limit some applications and the cell line can be difficult to transfect.

In this study, we conducted an integrated analysis of the LUHMES neuronal model system, encompassing genetic, epigenetic, and transcriptomic approaches. Our assessment revealed the temporal expression patterns of *UBE3A-ATS* and 11,834 transcripts genome-wide during differentiation of LUHMES to neurons. Furthermore, we demonstrate differential expression of multiple genes within the AS/PWS imprinted locus following neuronal differentiation and the distinct strand-specific expression profiles. Notably, we uncovered and validated two CTCF loop interactions unique to LUHMES neurons from *MAGEL2* to *SNRPN* and from *PWAR1* to *UBE3A.* These developmentally induced changes in chromatin architecture support the neuron-specific changes to parental differentially methylated regions associated with gene imprinting at this locus.

## Results

### *UBE3A-ATS* is progressively induced during neuronal differentiation of LUHMES

We hypothesized that LUHMES may be a particularly useful model for the complex developmental dynamics of the AS/PWS locus and sought to further characterize its morphological, genetic, transcriptional, and epigenetic characteristics. LUHMES cells showed an epithelial-like morphology in the undifferentiated state but demonstrate morphological characteristics of neurons including long neurites resembling mid-brain axonal networks within seven days in differentiation media (Figure 2).

**Figure 2.**
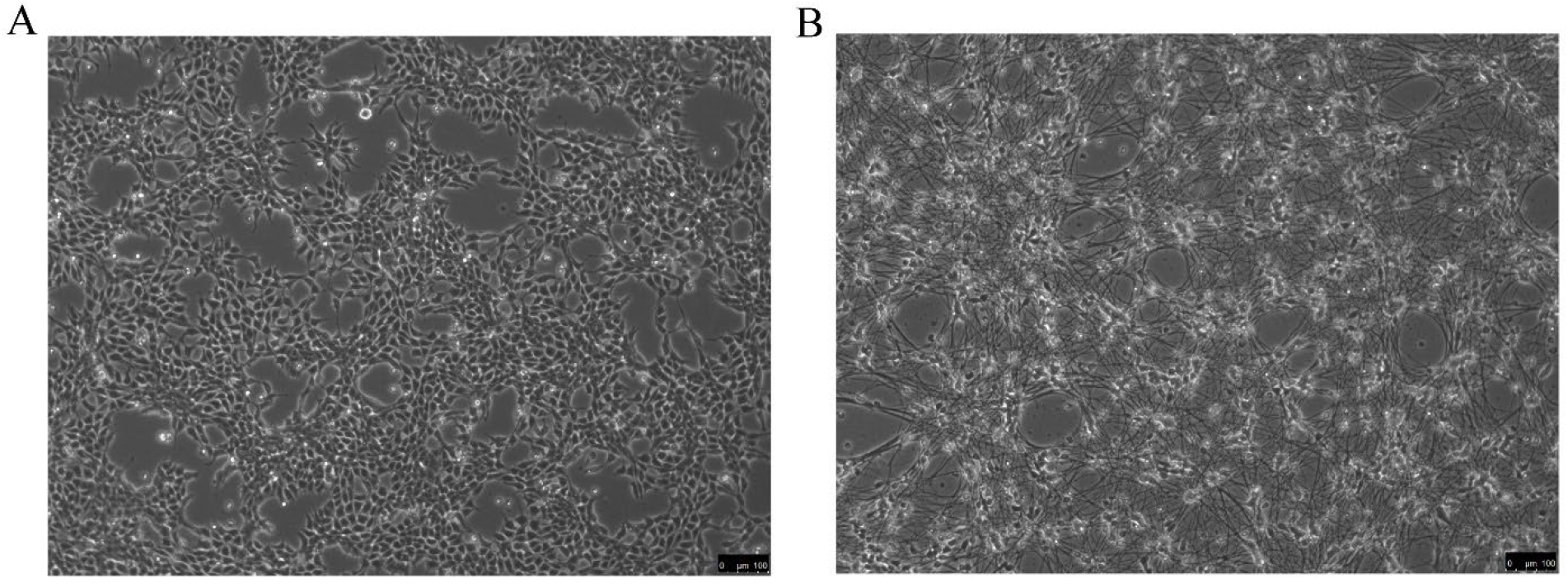
Brightfield Microscopy at 10X of **A.** Undifferentiated LUHMES. **B.** 7 day differentiated LUHMES neurons.

To evaluate the relevance of the LUHMES differentiation system for the postnatal neuronal dynamics of the *UBE3A* locus in AS, we evaluated the expression levels of the *UBE3A-ATS* transcript by quantitative reverse transcription PCR (qRT-PCR) across several cell types and human brain tissue. Controlling for input RNA and using PPIA as the housekeeping gene, we utilized the 2^-ΔΔCt^ method to calculate relative *UBE3A-ATS* transcript levels in HEK293T cells, undifferentiated LUHMES cells, 7-day differentiated LUHMES neurons and adult human cerebral cortex tissue (Figure 3). In the HEK293T cells and undifferentiated LUHMES cells, *UBE3A-ATS* transcripts were below the level of detection. In contrast, the differentiated LUHMES neurons showed high levels of *UBE3A-ATS*, which was comparable to that observed in adult cerebral cortex (Figure 3A). We then used qRT-PCR to characterize the temporal expression of the antisense transcript over a seven-day period in differentiation media (Figure 3B). On Day 1 and 2, *UBE3A-ATS* transcript levels were relatively low. However, by Day 4, there was a substantial increase in expression. This upward trend continued throughout the seven-day period, with the most substantial increases observed between Days 5 and 7. Together, these results demonstrate that LUHMES neurons are a valid model for the transcriptional changes in *UBE3A-ATS* expression known to occur during early postnatal neuronal maturation in the brain (26).

**Figure 3.**
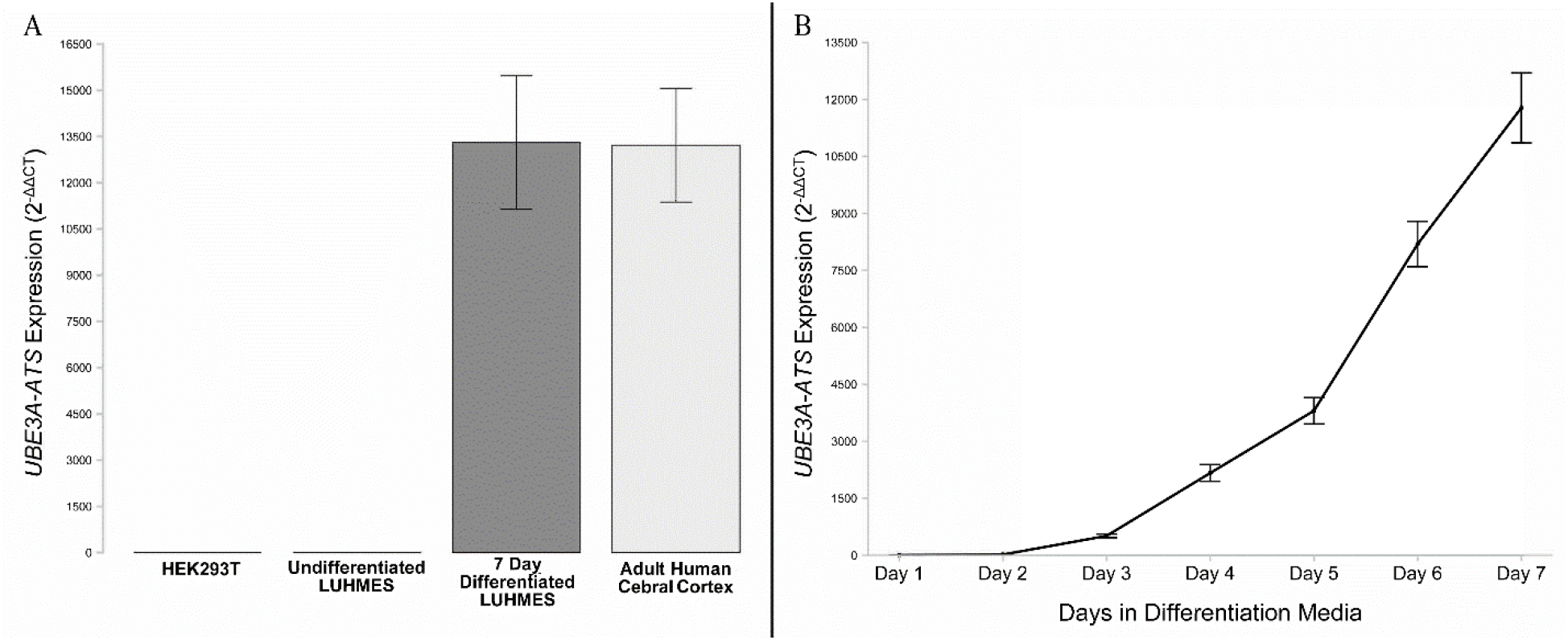
Quantitative RT-PCR. demonstrates that **A.** *UBE3A-ATS* is expressed in differentiated LUHMES neurons and human brain cortex, but not in undifferentiated LUHMES and HEK293T cells. **B.** LUHMES *UBE3A-ATS* transcript levels progressively increased throughout 7 days in neuronal differentiation media.

### Large-scale transcriptome changes, including 15q11-q13 imprinted genes are associated with LUHMES differentiation to neurons

To further characterize the transcriptional changes in 6-day differentiated LUHMES neurons compared to the undifferentiated state, we performed RNA-seq in triplicate cultures. When looking at differentially expressed genes in LUHMES neurons, we found 5,379 genes upregulated and 6,455 genes downregulated compared to undifferentiated LUHMES after correction for genome-wide significance (Figure 4**; S1A**). In LUHMES neurons, the top ten differentially expressed genes, based on the lowest adjusted P values, were *ALCAM, MAP2, RTN1, NCAM1, CNTN2, AKAP6, KIF5A, SCD5, ROBO2,* and *NRG1*. All these genes showed significant upregulation in LUHMES neurons compared to undifferentiated LUHMES cells, as indicated by log fold change (logFC) values ranging from 4.85 (*ROBO2*) to 8.66 (*CNTN2*). The adjusted P values for these genes ranged from 4.27E-13 to 8.98E-13, indicating significant differential expression (Figure 4A). The top ten differentially expressed genes that were downregulated in neurons were *H1-5, H2AC11, ASS1, H2BC18, NCAPD2, CCNB1, SMC4, SUSD2, HMGA2, and CENPF* with negative logFC values ranging from -4.56 (*NCAPD2*) to - 7.53 (*H1-5*). The adjusted P values for these genes ranged from 4.94E-13 to 1.72E-12, again indicating significant differential expression (Figure 4A).

**Figure 4:**
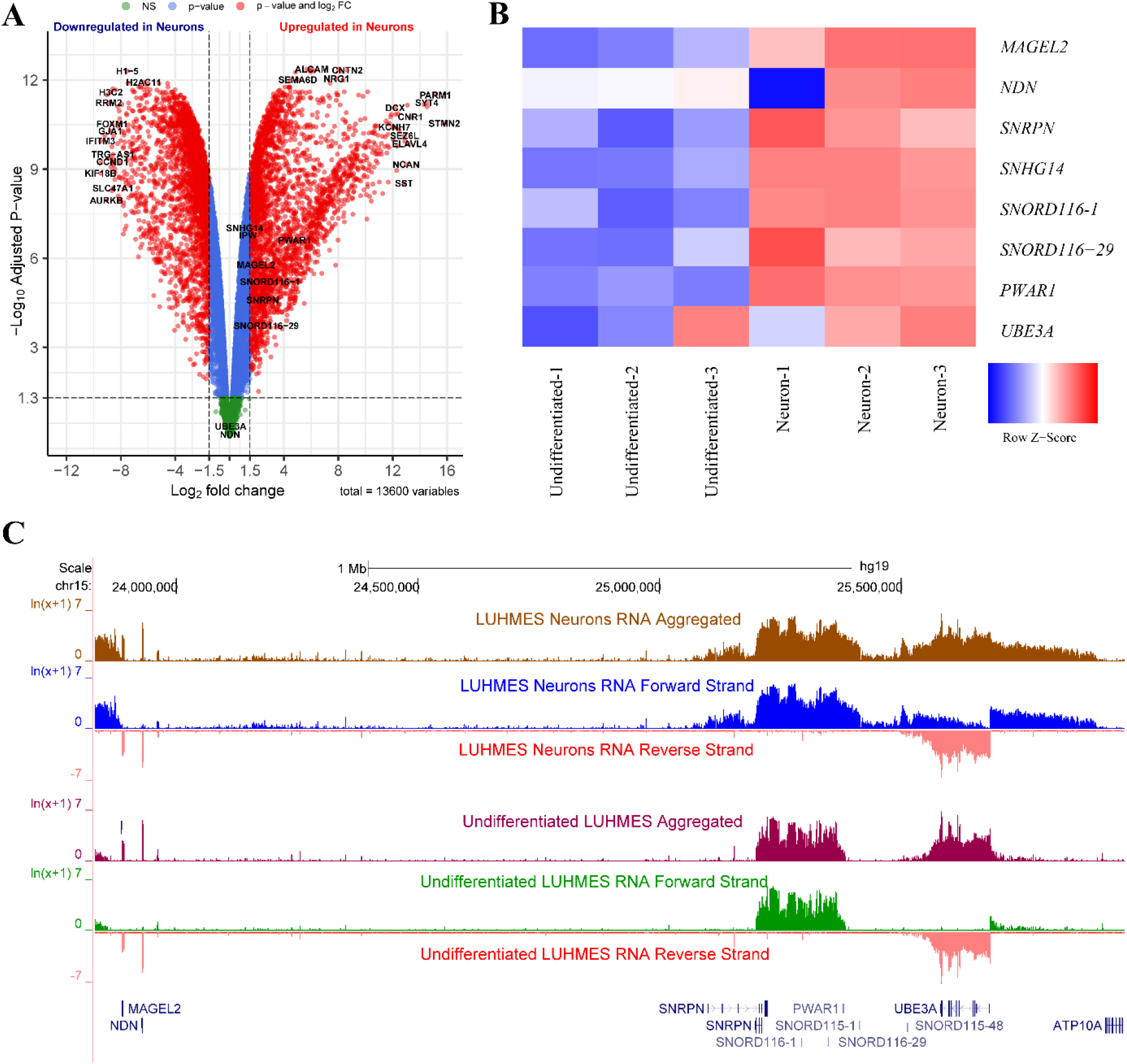
RNA-seq analyses. **A.** Volcano plot of differential gene expression comparing undifferentiated vs differentiated LUHMES in -Log_10_ P-value vs Log_2_ fold change. Green = Log_2_ fold change < |1.5|, P-value > 0.05, Blue = Log_2_ fold change < |1.5|, P-value < 0.05, Red = Log_2_ fold change > |1.5| and P-value < 0.05, those on the left upper quadrant with Log_2_ fold change < -1.5 represent genes downregulated in LUHMES neurons, while those on the right upper quadrant with Log_2_ fold change > 1.5 represent genes upregulated in LUHMES neurons. Genes of the 15q11-q13 and those with highest absolute values for Log_2_ fold change as well as smallest P-values are labeled in black. **B.** Heatmap of differentially expressed genes at the 15q11-q13 locus based on Z score, Red = Upregulated in Neurons, Blue = Downregulated, shown in triplicates (only first and last *SNORD116* are included). **C.** LUHMES aggregated and strand specific transcription from throughout the locus for Neurons and Undifferentiated cells (hg19: chr15:23,832,378-25,962,021).

Within the AS/PWS locus, several genes showed significant upregulation in LUHMES neurons compared to undifferentiated LUHMES cells. Notably, *MAGEL2, SNRPN, SNHG14, PWAR1,* and several small nucleolar RNAs (snoRNAs) within the *SNORD116* cluster showed significant upregulation, with logFC values ranging from 0.05 (*SNORD116-13*) to 5.20 (*SNORD116-24*). The *UBE3A* gene, which is of particular interest in the context of AS, showed a slight upregulation, but this was more variable and not statistically significant (logFC = 0.08, adjusted P value = 0.45) (Figure 4B).

In the visualization of strand-specific transcription, only the forward strand was distinctly altered between these two cell states (Figure 4C). In neurons, an increase in forward strand transcription was observed starting upstream of the PWS-ICR in the *SNRPN* 5’ alternative exons and extending past *UBE3A* in the antisense direction. In undifferentiated cells, however, the forward transcript started at the PWS-ICR within *SNRPN* and showed an abrupt decrease of forward strand transcription at the 3’ end of *PWAR1* as seen in previous studies (26) (Figure 4C). Based on the known parental expression of the genes in this locus, these results demonstrate that the elevated transcription following neuronal differentiation in LUHMES was specific to the paternal transcripts in the forward strand direction.

To identify the biological processes that were enriched in LUHMES neurons and undifferentiated LUHMES cells, we performed a gene ontology (GO) enrichment analysis (Figure 5). In LUHMES neurons, a reactome pathway analysis revealed enrichment for the dopamine neurotransmitter release cycle pathway (*p*=2.79E-09). This finding is consistent with the expected dopaminergic nature of LUHMES neurons and further supports their neuronal identity (Figure 5A). The GO cellular component showed that the top processes enriched in neurons are related to neuron projection (*p*=1.81E-7) and axonal development (*p*=7.23E-17) (Figure 5B). Additionally, the top five enriched biological processes were nervous system development (*p*=1.62E-17), axonogenesis (*p*=2.05E-15), synapse organization (*p*=3.08E-15), axon guidance (*p*=1.24E-13), and modulation of chemical synaptic transmission (*p*=1.00E-11) (Figure 5C). These processes are all critical for neuronal function and development, suggesting that the genes upregulated in LUHMES neurons are involved in these key biological processes. In contrast, the top five enriched biological processes downregulated in LUHMES neurons were ribosome biogenesis (*p*=2.37E-76), gene expression (*p*=1.19E-72), translation (*p*=2.47E-72), rRNA processing (*p*=3.93E-72), and cellular macromolecule biosynthetic process (*p*=1.75E-71) (Figure 5D). These processes are fundamental for cellular function and growth which non-dividing cells likely downregulate as growth slows and ceases.

**Figure 5:**
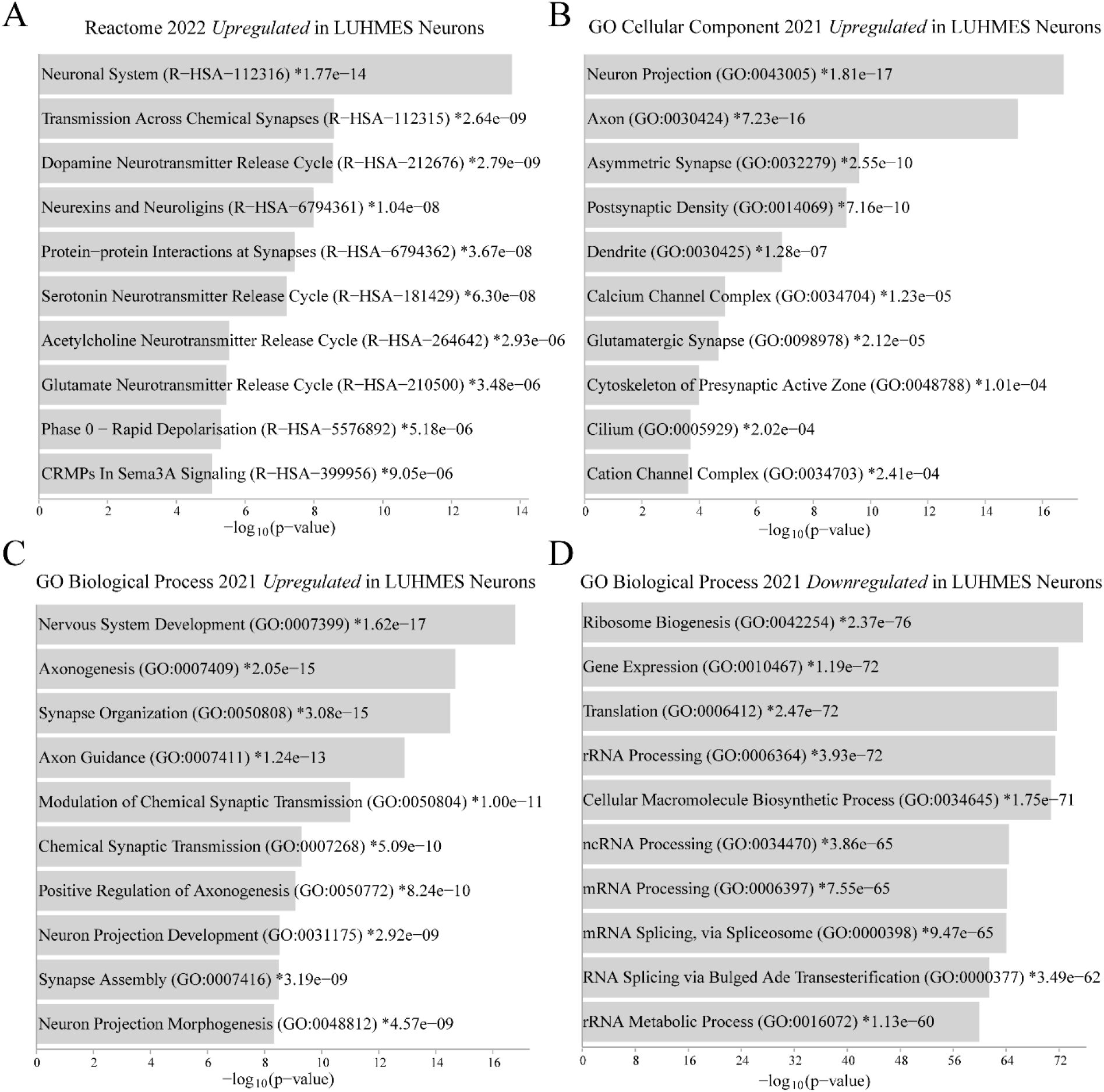
Enrichment analysis for genes upregulated and downregulated LUHMES neurons. **A.** Reactome 2022 GO terms enriched in LUHMES neurons **B.** GO cellular component 2021 enriched in LUHMES neurons. **C**. GO biological process 2021 for genes upregulated in LUHMES neurons. **D.** GO biological process 2021 for genes downregulated in LUHMES neurons.

### Chromatin loop analysis revealed neuron-specific CTCF loops in LUHMES

CTCF is a key regulator of chromatin architecture and its role in the formation of chromatin loops is crucial for gene regulation. We employed HiChIP analysis to investigate differential chromatin loop formations involving CTCF in undifferentiated and 6-day differentiated LUHMES neurons. Our approach utilized two protocols using the FitHiChIP pipeline: the first, a stringent analysis across all interactions at an FDR of 0.05, and the second, a looser criterion requiring peaks to be present in at least two replicates, set at a more permissive FDR of 0.1 (27). Significant interactions were detected by CHiCAGO with a score threshold of ≥5 (27,28). To understand the role of CTCF loop dynamics in the AS/PWS locus, we focused on a region spanning chr15:23,832,378-25,962,021 (hg19). Specifically in the differentiated LUHMES neurons, we observed a significant long-range chromatin loop interaction spanning approximately 1.2 Mb between the *MAGEL2* gene (chr15:23,890,148-23,895,147) and a region ∼100 kb upstream of *SNRPN* (chr15:25,090,148-25,095,147) (Figure 6). This interaction was given a confidence score of 6 by our stringent analysis. In comparison, *MAGEL2* also interacts with a cluster of loops present in both cell types with confidence scores that range from single digits to 169 (chr15:23,890,148-24,105,147). Another notable neuron specific chromatin loop interaction was observed between the *PWAR1* gene (chr15:25,380,148-25,385,147), and a region located approximately 64 kb downstream of the *UBE3A* (chr15:25,745,148-25,750,147) with a confidence score of 9 (Figure 6A). When using the less stringent filtering method, we also observed another interaction originating from the same *PWAR1* bin to the *UBE3A* promoter at (chr15:25,680,148-25,685,147) that was unique to neurons with a confidence score of 6 (Figure 6B). The *UBE3A* promoter bin also showed a nearby interaction within the 3’ *UBE3A* body (Figure 6B). Using this less stringent filtering method, we were able to identify 36,816 HiChIP interactions unique to neurons, 74,469 unique to undifferentiated cells and 26,162 shared between them (**Figure S1B**). We also observed some overall differences between the two cell types when looking at their contact matrix, with undifferentiated LUHMES showing a greater number of overall contacts (**Figure S2**).

**Figure 6:**
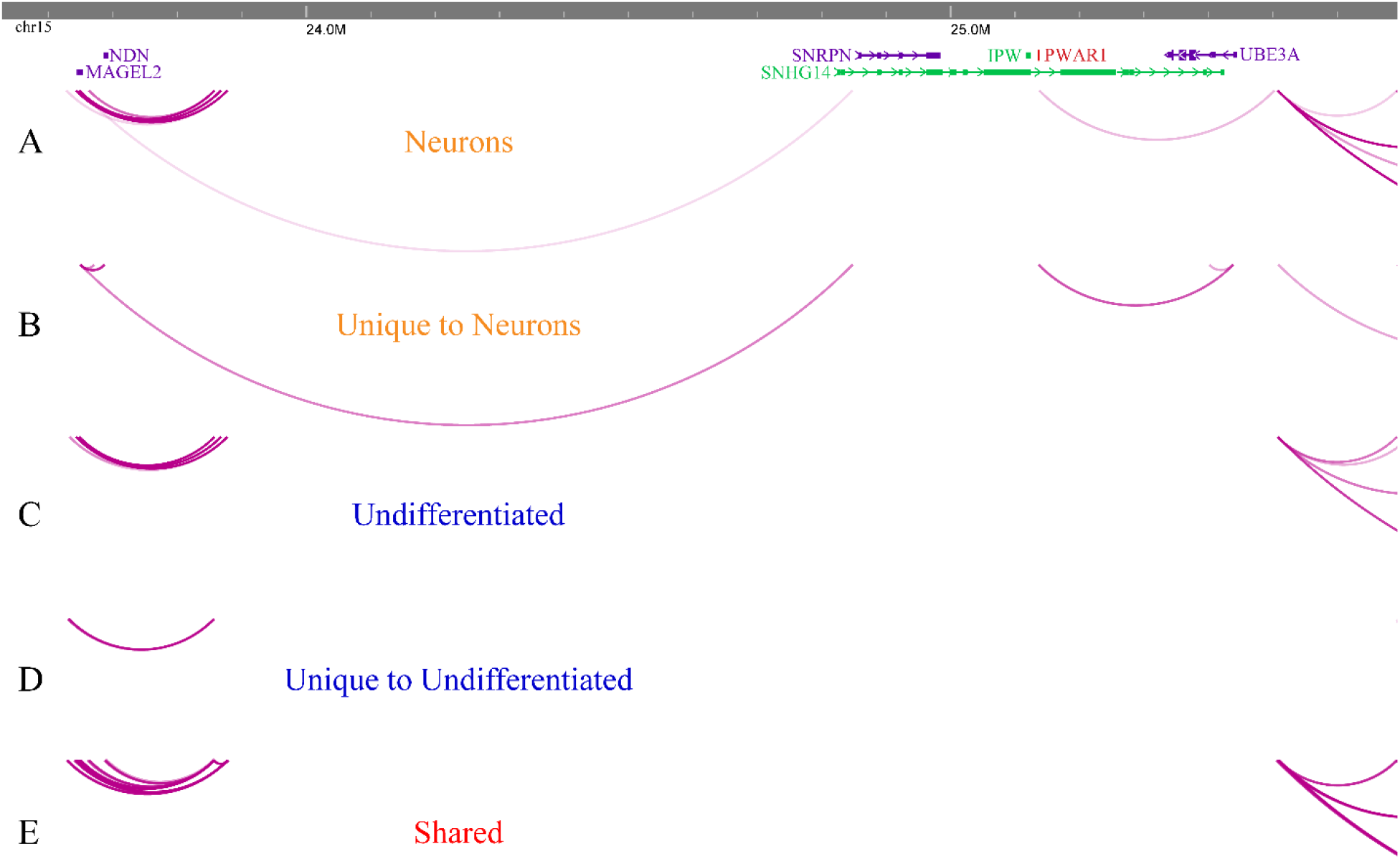
Neuron-specific long-range looping within the AS/PWS locus. WashU epigenome browser depiction of the CTCF HiChIP demonstrating long range chromatin interactions for LUHMES neurons and undifferentiated LUHMES. HiChIP loops are shown as purple arches, with greater intensity reflecting greater significance. **A.** Loops detected in neurons with stringent filtering using 5 kb bins at 0.05 FDR **B.** Loops unique to neurons using a less stringent analysis and FDR 0.1. **C.** Loops in undifferentiated cells using 5 kb bins at 0.05 FDR. **D.** Loops in undifferentiated LUHMES using less stringent analysis, 0.1 FDR. **E.** Only loops shared between the two cell types at 0.1 FDR.

### 4C validation of HiChIP findings

While HiChIP provides a non-biased all-to-all comparison of all interactions associated with CTCF genome-wide, circular chromosome conformation capture (4C) is a one-to-all comparison of genome-wide chromatin interactions with a specified genomic viewpoint. To validate the chromatin loops identified in our HiChIP analysis, we performed several 4C experiments using viewpoints from *SNRPN*, *PWAR1*, and *UBE3A* (Figure 7). A loop observed exclusively in neurons from *MAGEL2* to *SNRPN* in the HiChIP data was replicated by an interaction from the *SNRPN* viewpoint (chr15:25,092,529) to a region which included *MAGEL2* (chr15:23,885,650-23,902,345) (Figure 7A). In contrast, from the same viewpoint in undifferentiated LUHMES, a different loop interaction was detected spanning 97,966 bp (chr15:25,318,254-25,332,219) (Figure 7B). Upon examining the neurons from the *PWAR1* viewpoint (chr15:25,382,560), we observed two nearby interactions between the 5’ end of the *SNORD115* cluster (chr15:25,411,021-25,514,112) and a region encompassing *UBE3A* (chr15:25,514,112-25,689,001), validating the *PWAR1*-*UBE3A* interaction we saw using the less stringent HiChIP analysis (Figure 7C). In contrast, this *PWAR1*-*UBE3A* interaction was not present in undifferentiated LUHMES (Figure 7D), demonstrating specificity for neurons. Using a viewpoint centered on the *UBE3A* promoter (chr15:25,684,119), we confirmed that *UBE3A* interacted with the *PWAR1* region from chr15:25,371,923-25,384,018 (Figure 7E). In contrast, the undifferentiated LUHMES with the same viewpoint interacted with three entirely different regions forming three loops between the *SNORD115* cluster (chr15:25,549,549-25,561,569), the 5’ end of *UBE3A* (chr15:25,562,286-25,583,229), and a third downstream region close to *ATP10A* from chr15: 25907922-25926082 (Figure 7F).

**Figure 7:**
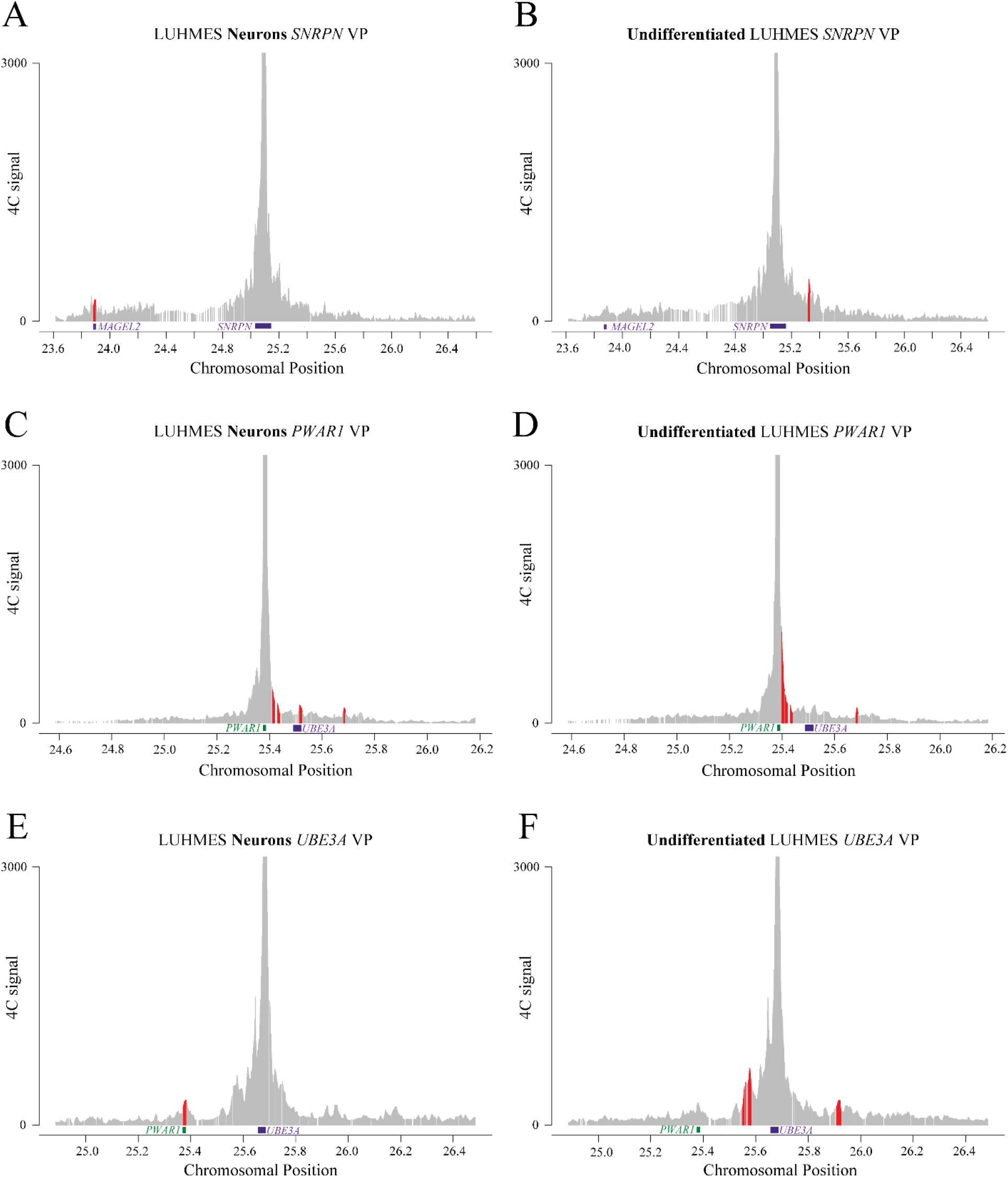
4C validates the HiChIP loop anchors differential in neurons. Results from a 4C analysis, in red are called interaction peaks (hg19) **A.** LUHMES neurons using *SNRPN* viewpoint (chr15:25092529.) **B.** Undifferentiated LUHMES using the same SNRPN viewpoint. **C.** Neurons using the *PWAR1* viewpoint (chr15:25382560). **D.** Undifferentiated using the same *PWAR1* viewpoint. **E.** LUHMES neurons using a viewpoint located at the promoter of *UBE3A* (chr15: 25684119). **F.** Undifferentiated LUHMES using the same *UBE3A* viewpoint.

### Integration of allele specific CpG methylation with CTCF loops and imprinted expression

We used Oxford Nanopore Technology (ONT) sequencing to examine CpG methylation differences within the AS/PWS locus, with a particular focus on the CTCF motif sequences where unique loops were found in neurons. This analysis provides us with insights into the relationship between DNA methylation, CTCF loops and gene expression that could potentially regulate imprinting of *UBE3A* and other genes within the AS/PWS. We used ONT’s pipeline, modkit pileup, to call methylation and visualize the data (**Figure S3**). Global methylation landscape patterns between the undifferentiated LUHMES and neurons were overall similar.

ONT’s long reads provided the advantage of allowing phasing of the methylome using nanomethphase (29), which is of particular importance for imprinted loci. UCSC browser track hubs were created for visualization together with our LUHMES HiChIP and RNAseq data as well as other genome annotations (Figures 8**, 9, S3, S4**). We were able to assign parentage for each haplotype based on the well characterized paternal hypomethylation of the PWS-ICR (30,31)(Figure 8B).

**Figure 8:**
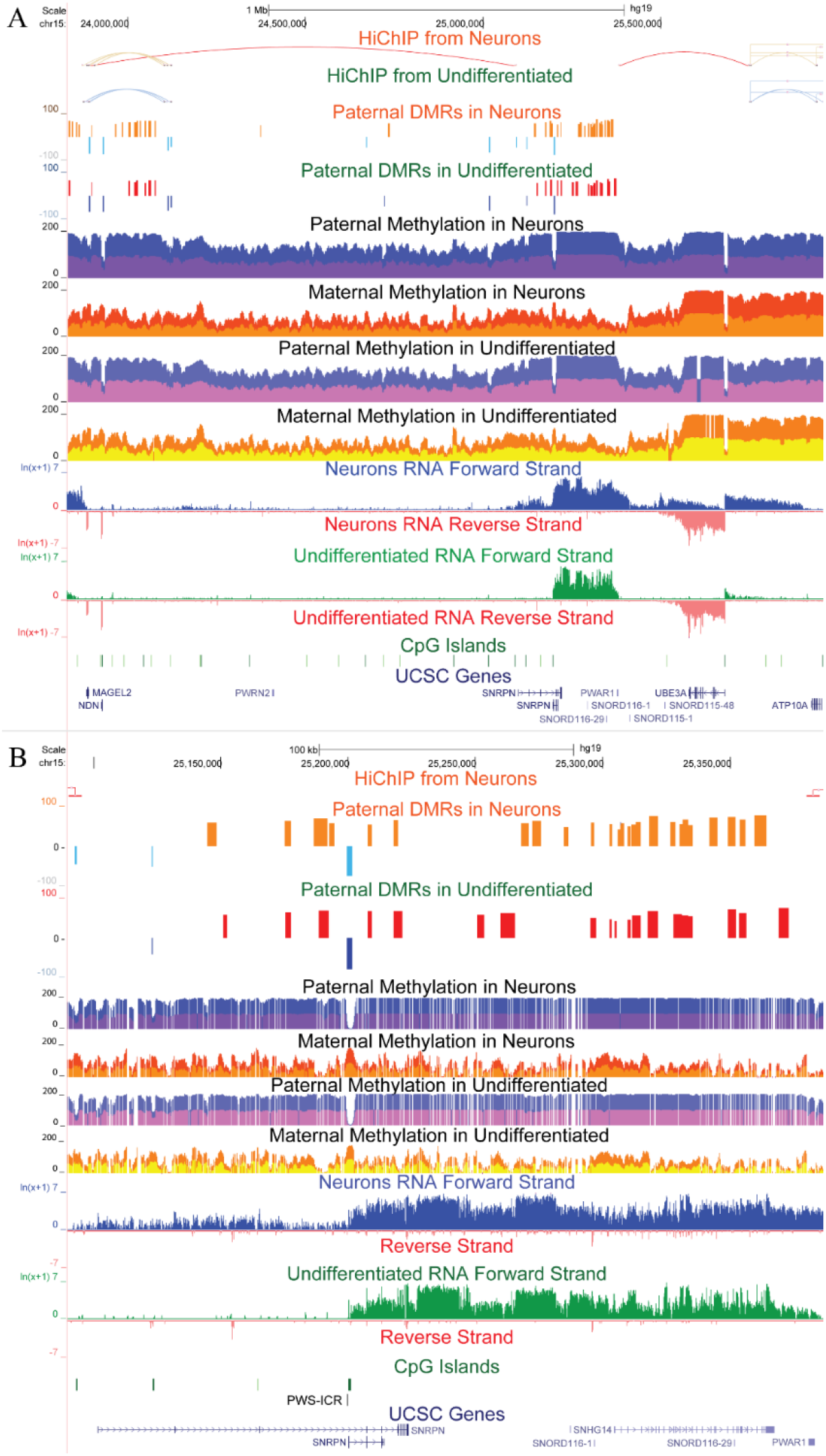
Integrated CTCF loops, paternal DMRs, methylation profile and strand specific transcription. Taken from the same LUHMES neurons and undifferentiated LUHMES samples. For neurons, paternally hypermethylated DMRs are shown in orange and paternally hypomethylated DMRs in light blue. In undifferentiated LUHMES, paternally hypermethylated DMRs are shown in red with dark blue representing hypomethylation. DMR values represent differences in percent methylation between paternal and maternal alleles. For methylation profiles 2 replicates were stacked and shown in different colors for contrast with values representing the sum of their percent methylation (max 200); CpG island and UCSC Genes are also included **A.** 15q11-q13 locus (hg19; chr15:23,832,378-25,962,021) **B.** A closer view of the paternal DMR cluster between *MAGEL2-SNRPN* and *PWAR1-UBE3A* neuron-specific loop anchors (chr15:25,089,681-25,387,210)

A distinct pattern of CpG methylation was observed in an allele-specific manner in both neurons and undifferentiated LUHMES cells (Figure 8A). We identified paternal-specific differentially methylated regions (DMRs), shown as tracks in Figure 8**-9** (blue, hypomethylated; red/orange, hypermethylated). Narrow regions of paternal hypomethylation were seen within broader regions of paternal hypermethylation, a pattern that was previously observed by whole genome bisulfite sequencing in postmortem PWS, AS, and Dup15q brain samples (32). When comparing the paternal allele to the maternal allele, the region between both neuron-specific chromatin loops was particularly enriched for paternal DMRs and was predominantly hypermethylated (Figure 8).

**Figure 9:**
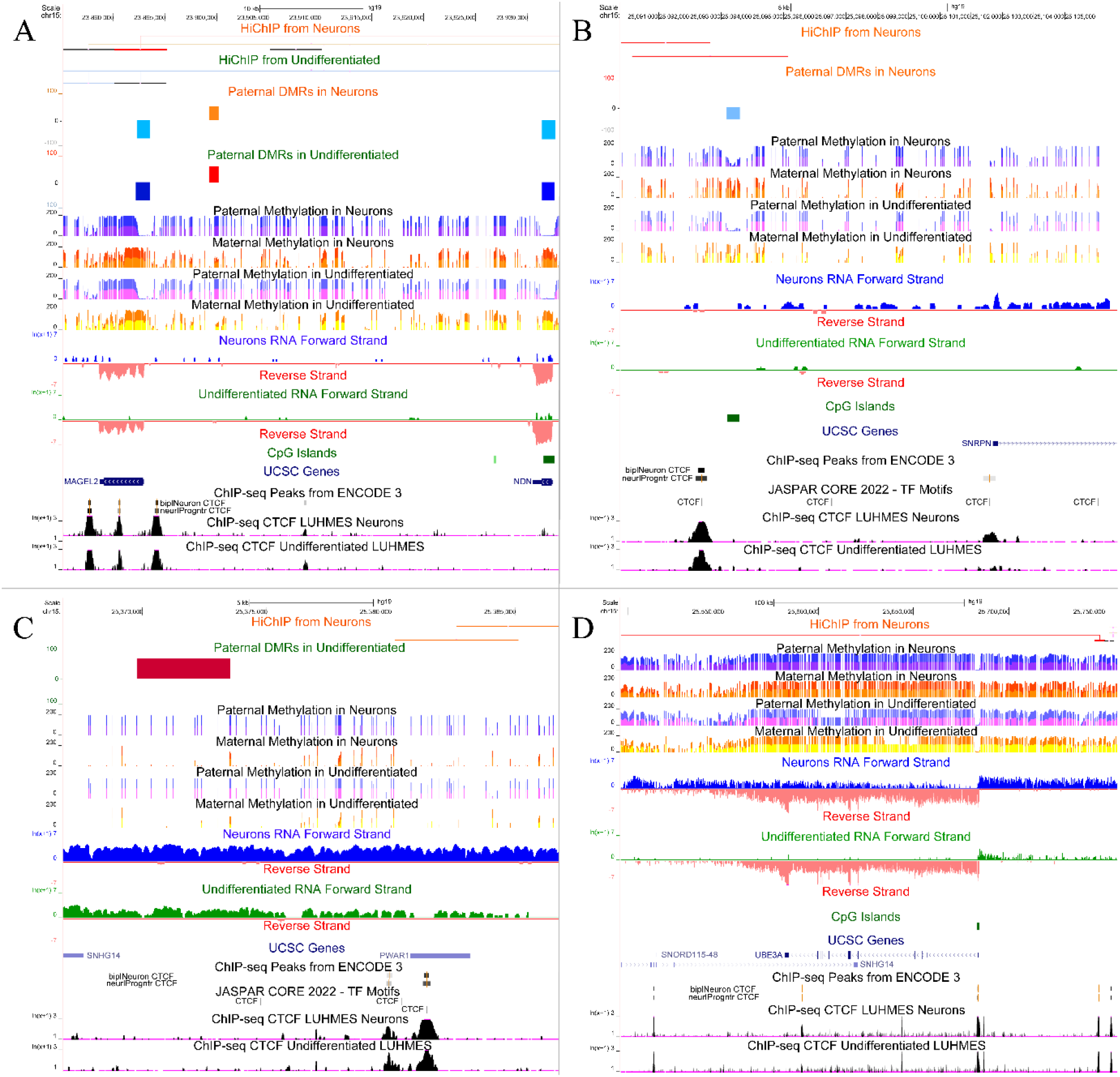
CTCF loops, paternal DMRs, methylation profile, transcription and ChIP-seq for each neuron specific loop anchor region. HiChIP and DMR tracks with no data in the region shown have been omitted. Additional tracks: ChIP-seq Peaks from ENCODE 3 for CTCF in bipolar neuron and neuronal progenitor were included to approximate LUHMES neurons and undifferentiated LUHMES respectively; JASPAR Core 2022 transcription factor motif for CTCF; LUHMES CTCF ChIP-seq from Pierce et al., 2018. **A.** A closer look at the region encompassing the *MAGEL2* anchor and *NDN* overlapping paternally hypomethylated DMRs (hg19; chr15:23,885,189-23,933,105) B. *SNRPN* upstream exon anchor that overlaps with a neuron-specific paternally hypomethylated DMR (chr15:25,089,792-25,105,663) C. *PWAR1* loop anchor and the paternally hypermethylated DMR exclusive to undifferentiated cells (chr15:25,366,804-25,386,789) D. *UBE3A* anchor region showing a hypermethylated profile but no paternal DMRs in either cell type (chr15:25,496,109-25,756,788).

For both cell states we observed paternally hypomethylated DMRs overlapping with the *MAGEL2* and *NDN* promoters with a region of hypermethylation in between (Figure 9A). However, a paternally hypomethylated region overlapping the *SNRPN* loop anchor was exclusive to neurons and corresponded to the start of *SNRPN* transcription specifically in neurons (Figure 8B**, 9B**). In contrast, a downstream paternal hypomethylated DMR at the PWS-ICR was associated with the beginning of *SNRPN* transcription in undifferentiated cells, despite its presence in both cell states (Figure 8B). A paternally hypermethylated DMR exclusive to undifferentiated LUHMES was observed upstream of the *PWAR1* anchor after which transcription decreases specifically in undifferentiated cells (Figure 9C). All loop anchors were adjacent to forward strand transcripts that increased expression in neurons (Figures 4**, 8, 9**). The loop anchor located nearest to *UBE3A* was about 61 kb 5’ of its biallelically hypomethylated CpG island promoter. A differential methylation analysis was also performed comparing the paternal allele in neurons to the paternal allele in undifferentiated cells, however this resulted in few additional DMR occurrences throughout a large portion of chromosome 15 (**Figure S4**). This was also true when comparing the maternal allele in neurons to the maternal allele in undifferentiated cells.

These findings provide novel insight into the dynamic changes in chromatin architecture that occur in the AS/PWS locus during the differentiation of LUHMES cells into neurons. The neuron-specific chromatin loops coincide with increased expression of multiple paternal transcripts, including *MAGEL2, NDN, SNHG14*, *SNRPN, PWAR1*, and *UBE3A-ATS* (Figures 4**, 8, 9**). These dynamic changes in neuronal chromatin structure associating with paternally expressed transcripts suggest their involvement in paternal silencing of *UBE3A*, although these experiments do not directly determine the allele-specificity of CTCF binding.

## Discussion

In this comprehensive study, we characterized and integrated genome-wide DNA methylation with CTCF loops and RNA expression of LUHMES cells to shed light on their relationship, with a particular emphasis on evaluating their potential as a model for AS. These results provide an integrative multi-omic atlas of neuronal differentiation in LUHMES that could be useful for investigations of multiple neurodevelopmental disorders. With particular relevance to PWS and AS, we identified two neuron-specific CTCF loop interactions: from *MAGEL2* to *SNRPN* and from *PWAR1* to *UBE3A*. A hypomethylated paternal DMR at the 5’UTR *SNRPN* anchor that corresponded to increased transcription of the paternal forward strand was also exclusive to neurons. Additionally, we observed a hypermethylated paternal DMR near the *PWAR1* anchor only in undifferentiated cells, suggesting its potential role in the regulation of the transcription boundary in this region for non-neurons. Our findings provide a robust foundation for the use of LUHMES as a human neuronal model, especially for understanding the epigenetic dynamics of parentally imprinted loci.

In the differentiation of LUHMES cells to neurons, we observed a swift upregulation of genes, notably the paternal transcripts from *MAGEL2* to *UBE3A-ATS*. This rapid expression of *UBE3A-ATS*, achieved within seven days, underscores the potential of LUHMES cells as a superior model for neuronal studies, particularly over iPSC-derived neurons where differentiation is more protracted. The swift transition of LUHMES cells to mature neuronal functions, alongside their non-cancerous origin, offers a distinct advantage in delineating the boundary region of loci and assessing the regulatory role of CTCF in *UBE3A-ATS* expression. This expeditious differentiation, coupled with a downregulation of genes involved in broad cellular processes such as transcription and biogenesis, reflects a shift from cellular proliferation to specialization, positing LUHMES cells as a valuable tool for rapid and efficient neurogenetic investigations.

Within the PWS/AS locus, only the forward strand transcriptional profile was distinct between differentiated and undifferentiated LUHMES. All known forward strand transcripts within this locus are exclusively expressed from the paternal allele. In neurons we saw an increase in paternal forward strand transcription begin at the *SNRPN* upstream alternative exons and continue through and beyond the *UBE3A* gene body. The *UBE3A* gene body was also highly methylated on both alleles, in contrast to the broad swaths of maternal hypomethylation over the entire imprinted locus, consistent with previous results observed by Illumina methylome sequencing in PWS, AS, and Dup15q syndrome brain (32). In contrast, there was an abrupt decrease in forward strand transcription after *PWAR1* in undifferentiated cells, consistent with the prior finding of *PWAR1* as a transcriptional boundary in non-neurons (16).

While it has been known for decades that loss of the PWS-ICR affected not only transcript in *cis* starting from *SNRPN*, but also *MAGEL2* and *NDN* located ∼1 Mb away, the mechanism was not known (33). Here, we identified a neuron specific CTCF loop between *MAGEL2* and the alternative 5’ transcriptional start site of *SNRPN* which shows highest expression in brain and ovary (33). While the *MAGEL2*-*SNRPN* loop was observed less frequently than nearby interactions, it persisted with stringent filtering and was replicated by 4C from a separate LUHMES culture. The downstream anchor of the neuron-specific *MAGEL2*-*SNRPN* loop corresponded with both neuron-specific paternal loss of methylation and elevated paternal forward strand transcription, extending almost 2 Mb from *MAGEL2* through the distal side of the *UBE3A* gene body. While previous research has identified long-range chromatin loops emanating from the PWS-ICR using array-based methods, ours is the first sequencing-based analysis to characterize the specific interaction between *MAGEL2* and *SNRPN* in neurons (34). This chromatin loop interaction within the 15q11-q13 region enriches our understanding of the broad-scale architecture and epigenetic landscape, which is pertinent to the study of imprinting-related neurodevelopmental disorders linked to this segment of the genome.

Interestingly, the DMR closest to the *PWAR1* loop anchor was uniquely paternally hypermethylated in undifferentiated LUHMES. Given that adjacent DNA hypermethylation is known to hinder CTCF binding, this observation may relate to mechanisms similar to those at the *Igf2* locus in mice. There, hypermethylation near the *Igf2* gene prevents CTCF from binding to its regulatory sequences, which in turn affects the gene’s expression (35,36). Both the hypomethylation near *SNRPN* in neurons and the hypermethylation at *PWAR1* in undifferentiated LUHMES are states that favor CTCF binding and subsequent loop configuration observed in the differentiated state. Such dynamic DNA methylation and chromatin loop interactions could represent the necessary and/or sufficient conditions for favoring transcriptional progression past *PWAR1* and through *UBE3A-ATS*, potentially silencing the paternal *UBE3A* allele in neurons. It is noteworthy that induced DNA demethylation has been previously employed to reshape chromatin topology at the *IGF2-H19* locus (37).

One of the prevailing models describing how *UBE3A* is silenced by *UBE3A-ATS* within the AS/PWS locus is the collision model, which proposes that RNA polymerase II from convergent transcripts can disrupt gene expression (23,24). Our analyses suggest a novel view of how the *SNHG14* lncRNA, beginning at the PWS-ICR, could regulate *UBE3A-ATS* expression to silence paternal *UBE3A* in neurons (Figure 10). A chromatin interaction between *MAGEL2* and *SNRPN*, either alone or in combination with the loop from *PWAR1* to *UBE3A*, could be enhancing forward strand transcriptional progression through the *UBE3A* gene body specifically in neurons, thereby upsetting the balanced transcriptional collision seen at *PWAR1* in non-neurons.

**Figure 10.**
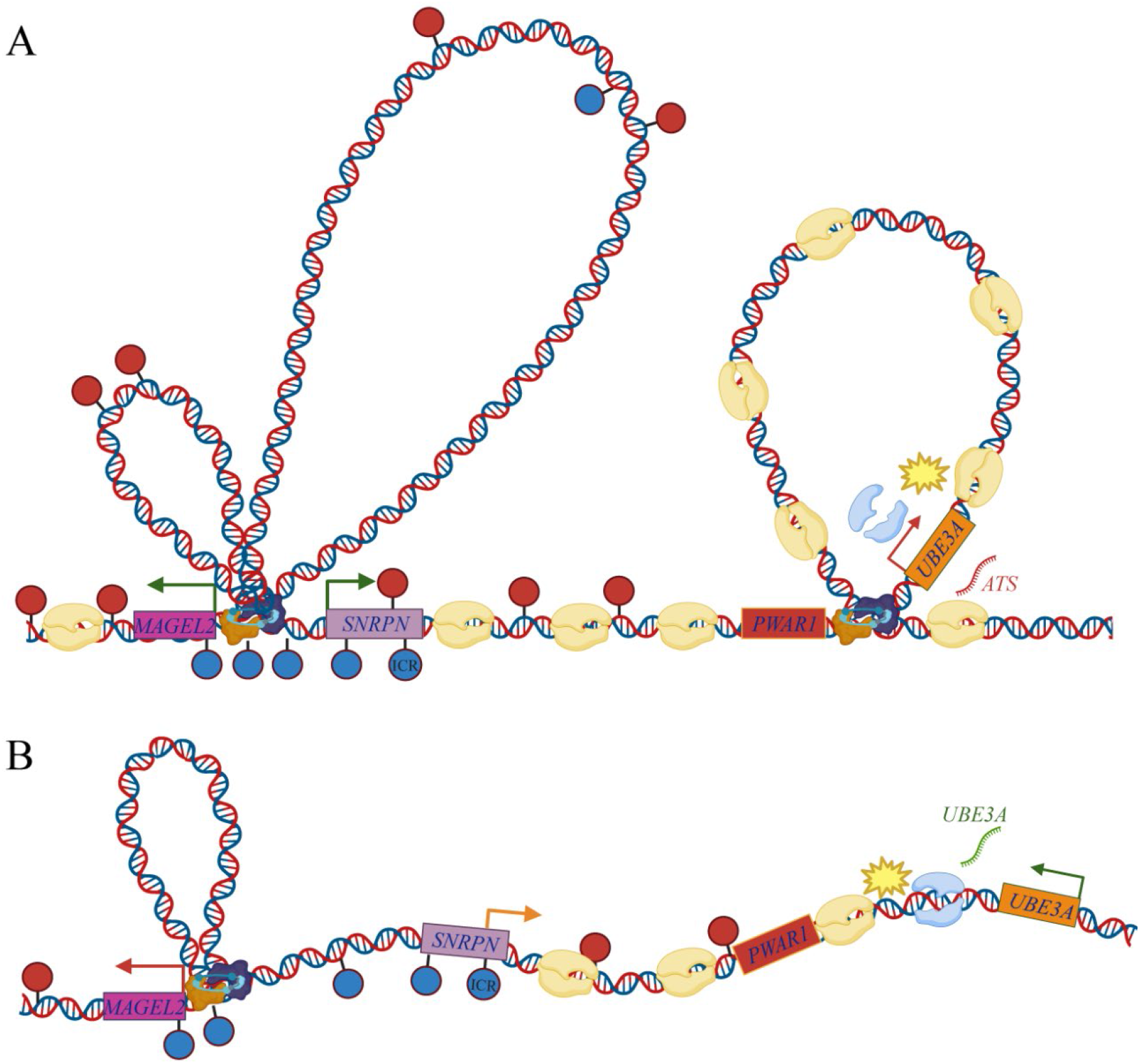
Proposed model of the epigenetic landscape on the paternal allele at the human 15q11-q13 locus in. **A.** LUHMES Neurons and **B.** Undifferentiated cells. Boxes represent labeled gene regions, arrows demonstrating direction of transcription with green representing transcribed genes, red representing silent genes and orange relatively reduced transcription. Paternal DNA hypermethylation is represented with red lollipops, and hypomethylated regions with blue lollipops. RNA polymerase II molecules are shown in yellow for forward strand transcription and blue for reverse strand transcription, while yellow star denoting proposed site of polymerase II collision. Note that the paternal hypermethylation close to *PWAR1* was only present in undifferentiated LUHMES. Not to scale, created with BioRender.com

To further investigate these mechanisms, ChIP-seq assays targeting Polymerase II, H3K27ac and transcription factors such as NRF1, TAF1, L3MBTL2, ZFX, and POLR2A could be insightful. These factors, involved in the enhancer interaction between *MAGEL2* and *NDN*, suggest a mechanism by which they could similarly influence the *MAGEL2-SNRPN* interaction (10,38).

The use of catalytically active CRISPR-Cas9 to remove CTCF binding sites, the use of artificial loops and the potential use of a split luciferase reporter system for visualization of loop manipulation in live cell models for neuronal differentiation represent additional innovative approaches at our disposal to investigate the regulation of neuronal *UBE3A* expression (39,40).

Building on these foundational insights, our research supports the use of artificial transcription factors (ATFs) to modulate the epigenetic environment surrounding the *UBE3A* gene. Consisting of a catalytically inactive Cas (dCas) fused to an effector domain, ATFs can be designed for specific epigenetic modifications, either to demethylate or methylate DNA at targeted sites (41,42).

Given our capability to edit DNA methylation in the mammalian genome, we envision employing dCas-TET1 for demethylation at the *PWAR1* binding site to explore its role in halting lncRNA expression past this region in non-neuronal cells (43). Furthermore, the use of dCas-DNMT3AL to hypermethylate the *PWAR1* binding site, alone or in conjunction with the *SNRPN* CTCF binding site in neurons, aims to replicate the transcriptional boundary characteristic of undifferentiated cells, providing insights into *UBE3A* silencing mechanisms. The persistence of epigenetic memory may require not only DNA methyltransferases like DNMT3AL but also histone methyltransferases such as Ezh2 or KRAB (44). Alongside DNA methylation, the capacity to edit histone states, including the recent applications of dCas9-HDAC, demonstrate one of the many potential avenues, further expanding our capacity to dissect and manipulate the epigenetic landscape (45,46).

The LUHMES cell model, with its human derivation and capability for rapid neuronal differentiation, is pivotal for our proposed experiments. It not only validates the biological relevance of our findings but also positions itself as a promising platform for bridging laboratory discoveries to therapeutic applications. This human-based system enhances the translational potential of our work, suggesting a clear path for the transferability of our findings to therapeutic interventions.

Our proposed strategy emphasizes the strategic importance of leveraging targeted epigenetic modifications within the LUHMES model to explore complex regulatory mechanisms of *UBE3A*. The advancements of epigenetic editing into clinical trials underscore the timeliness of our approach (46). While our heterozygous SNP data can assist in allele-specific editing, it is not a necessity for the creation of maternal *UBE3A* knockouts, which can be verified by assessing its expression in the undifferentiated state and its absence in differentiated LUHMES neurons. By highlighting the feasibility of translating our insights into clinical applications, we underscore the potential of our research in alleviating symptoms for patients living with AS and other neurogenetic disorders with an epigenetic component, marking a significant step toward innovative therapeutic solutions.

## Materials and Methods

### Cell culture

LUHMES were purchased from ATCC catalog number CRL2927 (2021). They were cultured and differentiated as described in Scholz *et al* 2011 with minor optimizations. We found that only using hydrophilic Nunclon® Δ surface treated flask allowed for appropriate cell adherence (Sigma, Cat. No. F7552-1CS. We also modified the additional coating of poly-L-ornithine (Sigma, Cat. No. P-3655-10MG) and fibronectin (MilliporeSigma, Cat. No. 341631-1MG) by mixing 50 ug/mL of poly-L-ornithine and 1 ug/mL of fibronectin, covering the flask and incubating overnight at 37 C. We found that two rinses with water was enough for the subsequent wash. For maintenance media we used DMEM/F-12, GlutaMAX™ (Gibco-Invitrogen, Cat. No. 10565-018) with 1% N2 supplement (Gibco-Invitrogen, Cat. No. 17502-048). Undifferentiated LUHMES were cultured to 80% confluency by removing media, washing with Ca++/Mg++ free Dulbecco’s phosphate-buffered saline (D-PBS) and incubating in warm 0.025% trypsin in D-PBS at 37 C for 2 minutes followed by light scraping. For differentiation we used the maintenance media with the addition of 2 ng/ml of human recombinant gDNF (Thermo, Cat. No. PHC7045), 1 mM of dibutryl cAMP (Sigma, Cat. No. D0627) and 1 ug/ml of tetracycline (Sigma, Cat. No. T7660-5G). Cells were place in differentiation media to 50-70% confluency and the first day was considered day 0. Differentiation media was changed every other day while leaving approximately 20% of the prior media. Since LUHMES neurons may become detached on day 7 those cells were harvested on day 6. Differentiated LUHMES were harvested in the same manner as undifferentiated LUHMES. Technical replicates used for the HiChIP, RNA-seq and ONP were all grown from the same aliquot of frozen (passage 4) LUHMES. HEK 293T (CRL-3216) were purchased from ATCC and manufacturers recommendations were followed for growth and subculturing.

### qPCR

qPCR assays were carried out on a Bio-Rad CFX384 real time system. Thermo-Fisher’s TaqMan gene expression probes for *UBE3A-ATS* (FAM hs01372957_m1) were used with PPIA (VIC hs99999904) as the housekeeping gene. To calculate values the 2^-ΔΔCt^ method was used where ΔΔCt was calculated by subtracting ΔCt of control (*PPIA*) from ΔCt of experimental (*UBE3A-ATS*). Postmortem human cortex (#1406) was obtained from the Maryland Tissue Bank.

### RNA-seq

All replicates for RNA-seq were harvested from the same passage and time point as those used for the HiChIP, and ONP sequencing. RNA was isolated using Qiagen RNeasy Plus Mini Kit (Cat. No. 74134). To capture non-coding RNAs, expression was studied after ribosomal RNA depletion. Strand-specific and dual-barcode indexed RNA-seq libraries were generated from 450 ng total RNA each using the Kapa RNA-seq Hyper kit (Kapa Biosystems-Roche, Basel, Switzerland) and both the QIAseq FastSelect–5S/16S/23S ribodepletion and FastSelect rRNA Plant reagents (Qiagen, Hilden Germany) in combination, following the instructions of the manufacturers. The fragment size distribution of the libraries was verified in an automated electrophoresis platform on the TapeStation (Agilent, Santa Clara, CA). The libraries were quantified by fluorometry on a Qubit instrument (Life Technologies, Carlsbad, CA) and pooled in equimolar ratios. The pool was quantified by qPCR with a Kapa Library Quant kit (Kapa Biosystems) and sequenced on an Illumina NovaSeq 6000 (Illumina, San Diego, CA) with paired-end 150 bp reads. Results were processed using Babraham Bioinformatics TrimGalore (v0.6.7), STAR (2.7.3), Samtools (v1.17), and MultiQC (v1.9) using the GTF annotation file GRCh38.109 (47–49). Differential gene expression and visualization was performed in R with edgeR (v3.40.2), Lima-Voom (v3.54.2) and EnhancedVolcano (v1.16.0) in R (v4.2.1) (50–52). The overlapping region in Venn diagram (**Figure S1**) was calculated by subtracting all genes with an FDR < 0.05 from all genes with at least one read count in all samples. All differentially expressed genes with an FDR < 0.05 were used as input to https://maayanlab.cloud/Enrichr/ for GO enrichment analysis. For visualization in the UCSC Genome Browser pileup BAM files were concatenated from all replicates for each condition. CrossMap (v0.6.4) was used to lift over coordinates to hg19 using UCSC chain files (53). Strands were split, values were transformed by LOG (ln(1+x)) and we used a max range of 7 to highlight smaller peaks.

### HiChIP

Technical replicates of LUHMES neurons and their progenitors were collected and immediately flash-frozen. Chromatin was fixed with disuccinimidyl glutarate (DSG) and formaldehyde in the nucleus. Fixed chromatin was digested *in situ* with micrococcal nuclease (MNase) and then extracted upon cell lysis. Chromatin fragments were incubated with the CTCF antibody overnight for chromatin immunoprecipitation. The antibody-protein-DNA complex was pulled down with protein A/G-coated beads. Chromatin ends were repaired and ligated to a biotinylated bridge adapter followed by proximity ligation of adapter-containing ends. After proximity ligation, crosslinks were reversed, the DNA was purified from proteins and converted into a sequencing library. The sequencing library was generated using Illumina-compatible adapters.

Biotin-containing fragments were isolated using streptavidin beads before PCR enrichment of the library and the samples were run on a Novaseq 6000. The FitHiChIP (v9.1) pipeline was used to analyze the data using 5 kb bins with the parameters set to peak-to-peak or “stringent” with an FDR:0.05 and merged nearby loops (27). To identify loop interactions unique to a single condition, data were also analyzed using a less exclusive modified workflow where GenomicRanges (v1.48) and MACS2 (v2.2.9.1) were used to first filter out only the peaks present in two or more replicates before running through FitHiChIP with the parameters set to all to all or “loose” with an FDR:0.1 and merged nearby loops (54,55). Loops were categorized as shared or unique using BEDtools (v2.28) pairToPair with -type both imposed to ensure that both loop anchors were shared (56). Reads were initially mapped to GRCh38 and long-range interaction files were plotted in the WashU epigenome browser. To allow viewing of loops in the UCSC genome browser together with previous HiChIP assays, JASPAR scores and other useful tracks, coordinates were lifted to hg19 using the liftOver utility.

### 4C

We used three technical replicates for each viewpoint (VP) and condition harvested from a separate LUHMES cell thaw passage 4 with 10 million cells each. The 4C protocol was adapted from Krijger *et al* (2020) with the following modifications: Invitrogen MagMax DNAbinding beads (Cat. No. 4489112) were substituted for the Nucleomag beads (57). Primary restriction enzyme digest was performed using DpnII (Cat. No. R0543S) and secondary digestion with CviQI (Cat. No. R0639S) from NEB. Before sequencing a final cleanup using SPRI select beads from Beckman Coulter (Cat No. B23317) were used. The fragment size distribution of the library pool was verified via micro-capillary gel electrophoresis on a Bioanalyzer 2100 (Agilent, Santa Clara, CA). The pool was bead cleaned twice to remove the adapter-dimer at 129 bp. Then, the library was quantified by qPCR with a Kapa Library Quant kit (Kapa Biosystems/Roche, Basel, Switzerland). The library was sequenced on one flow cell of Aviti sequencer (Element Biosciences, San Diego, CA) with single-end 150 bp reads. Pipe4C (v1.1) was used for the initial data analysis followed by peak calling with PeakC (v0.2) using the default settings aligned to hg19 (57–59). Viewpoint (VP) primers included: *SNRPN* VP reading primer (FP) 5’-TGTAATCCCAACACACTGG-3’ and non-reading primer (RP) 5’-TGTTGTCTCTCATTTTCCTCA-3’. For the *PWAR1* VP FP 5’-TCATAGCTGAAACCATGAGA-3’ and RP 5’-TAGACGAACATTGCTGTGAC-3’ were used. For the *UBE3A* viewpoint FP 5’-ACCATCTTGGGAGACACAC-3’ and RP 5’-TCCTCATCTTGGTGGTAAAG-3’ were utilized.

### Oxford Nanopore Sequencing

We used two technical replicates from passage 4 for each condition that were from the same harvest as the HiChIP and RNAseq. Flash frozen cultured cell pellets containing 5 million cells were used for high molecular weight genomic DNA (gDNA) isolation. Two ml of lysis buffer containing 100 mM NaCl, 10 mM Tris-HCl pH 8.0, 25 mM EDTA, 0.5% (w/v) SDS and 100µg/ml Proteinase K was added to the frozen cell pellet. Each reaction was incubated at room temperature for up to 18 hours to ensure that the lysate was homogenous. The lysate was then treated with 20µg/ml RNase A at 37^0^C for 30 minutes and cleaned with equal volumes of phenol/chloroform using phase lock gels (Quantabio Cat # 2302830). The DNA was precipitated by adding 0.4X volume of 5M ammonium acetate and 3X volume of ice-cold ethanol. The DNA pellet was washed twice with 70% ethanol and resuspended in an elution buffer (10mM Tris, pH 8.0). Purity of gDNA was accessed using NanoDrop ND-1000 spectrophotometer. DNA was quantified with Qbit 2.0 Fluorometer (Thermo Fisher Scientific, MA). Integrity of the HMW gDNA was verified on a Femto pulse system (Agilent Technologies, Santa Clara, CA) where majority of the DNA was observed in fragments above 100 Kb. Sequencing libraries were prepared from 1.5µg of high molecular weight gDNA using the ligation sequencing kit SQK-LSK114 (Oxford Nanopore Technologies, Oxford, UK) following instructions of the manufacturer with the exception of extended incubation times for DNA damage repair, end repair, ligation and bead elutions. 30 fmol of the final library was loaded on the PromethION flowcell R10.4.1 (Oxford Nanopore Technologies, Oxford, UK) and run was set up on a PromethION P24 device using MinKNOW 22.12.5. To improve the yield, the flow cell was washed with a flow cell wash kit EXP-WSH004 (Oxford Nanopore Technologies, Oxford, UK) at approximately 24 and 48 hrs. after the start of the run and the fresh library was loaded.

Basecalling was performed after the run using guppy 6.5.7. For the non-phased methylation data 5mc-5hmc calls were also made with guppy. The ONP pipeline’s modkit-pileup (v0.1.11) was employed using the --cpg option, with the GRCh38.p13 reference genome FASTA nucleic acid file. Subsequently, the UCSC liftOver tool was utilized to convert the coordinates to the hg19 reference genome. For the phased data, minimap2 (v2.24) was used for alignment to hg19 (60). f5c (v1.3) was used to call-methylation using the --pore r10 option (61). Clair3 (v1.0.4) was used to call variants using model r1041_e82_400bps_sup_g615 and whatshap (v2.0) was used for phasing (62,63). Nanomethphase (v1.2.0) was used to phase the methylome and DSS (v3.18) was used for differential methylation analysis (29,64–67). Paternal differentially methylation analysis was performed comparing paternal vs maternal, in both cell types using two replicates for each condition. Differentially methylated regions (DMRs) were called using the paternal allele as the treatment group and the maternal as the control from the same sample. As a result, positive values represent regions where methylation is higher in the paternal allele while negative represents higher methylation on the maternal allele, with those values indicating their percent differences. An additional analysis was performed comparing DMRs between neurons vs undifferentiated cells on the paternal allele and separately for the maternal allele (**Figure S3**). The bedGraphToBigWig was used to prepare visualization for UCSC Genome Browser (68). The well characterized hypomethylated region at the PWS-ICR was used to assign parentage for each haplotype.

## Acknowledgements

**Author contributions:** Osman Sharifi provided substantial bioinformatic assistance, critical advice and helped in 4C library clean up. Nicholas Heath contributed in the 4C design and experiment. Daniela Soto provided substantial bioinformatic support. J Antonio Gomez aided in LUHMES sample harvest. Dag Yasui contributed to the design of experiments, executed the isolation of high molecular weight DNA, assisted in the isolation of RNA for qRT-PCR and provided critical advice. Aron Mendiola contributed bioinformatically to the differential expression analysis. Henriette O’Geen, Ulrika Beitnere, Marketa Tomkova and Viktoria Haghani provided critical advice. Greg Dillon provided scientific input on the utility of the LUHMES model for preclinical testing. David J Segal and Janine M LaSalle conceived the study, contributed substantially on study design, data interpretation and the manuscript for this project.

The library preparation and sequencing for the Hi-ChIP analysis was carried out by Dovetail Genomics with Cory Padilla rendering invaluable assistance in the bioinformatic analysis. The library preparations and sequencing for all other assays were carried out at the UC Davis Genome Center DNA Technologies and Expression Analysis Core, supported by NIH Shared Instrumentation Grant 1S10OD010786-01, with notable contributions from Ruta Sahasrabudhe providing technical assistance in sample processing and sequencing, alongside advisory input.

The authors express their gratitude to Kyle Fink, Satoshi Namekawa, and Fred Chedin for their guidance, to Sophia Hakam, Madison Hypes, and He Yang for administrative support, and to Taylor M Koopot and Claudia Barquero Olivares for their assistance with the figures.

Data has been submitted to NCBI at http://www.ncbi.nlm.nih.gov/bioproject/1076600 Bioinformatic code used can be found at https://github.com/oran-gutierrez-fugon

## Abbreviations

AS: Angelman Syndrome
CTCF: CCCTC-Binding Factor
DMR: Differentially Methylated Regions
DSG: Disuccinimidyl Glutarate
D-PBS: Dulbecco’s Phosphate-Buffered Saline
FP: Forward Primer / Reading Primer
GEMs: Gel Bead-in-Emulsions
gDNF: Glial Cell-Derived Neurotrophic Factor
GO: Gene Ontology
HDAC: Histone deacetylases
ICR: Imprinting Control Region
iPSCs: Induced Pluripotent Stem Cells
logFC: Log Fold Change
lncRNA: Long Non-Coding RNA
LUHMES: Lund Human Mesencephalic
MNase: Micrococcal Nuclease
ONP: Oxford Nanopore
PWS: Prader Willy Syndrome
qRT-PCR: Quantitative Reverse Transcription Polymerase Chain Reaction
RP: Reverse Primer / Non-Reading Primer
snoRNA: Small Nucleolar RNAs
*UBE3A-ATS*: *UBE3A* Antisense Transcript
VP: Viewpoint

