## Supplemental Figures for "Integration of CTCF Loops, Methylome, and Transcriptome in Differentiating LUHMES as a Model for Imprinting Dynamics of the 15q11-q13 Locus in Human Neurons"

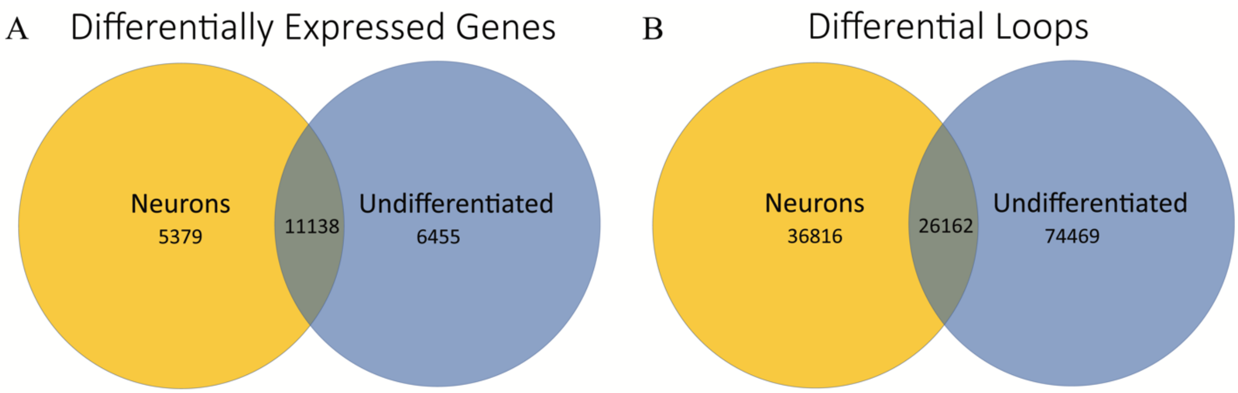


**Figure S1: Genome wide differential expression and differential loops. A.** Genes expressing differential expression FDR < 0.05 in Neurons (5,379) and undifferentiated LUHMES (6,455), overlapping region (11,138) was calculated by subtracting all genes with an FDR < 0.05 from the total gene pool that had at least one read count in any sample (22,972). **B.** Differential loops determined by the lower stringency analysis method described in HiChIP methods.

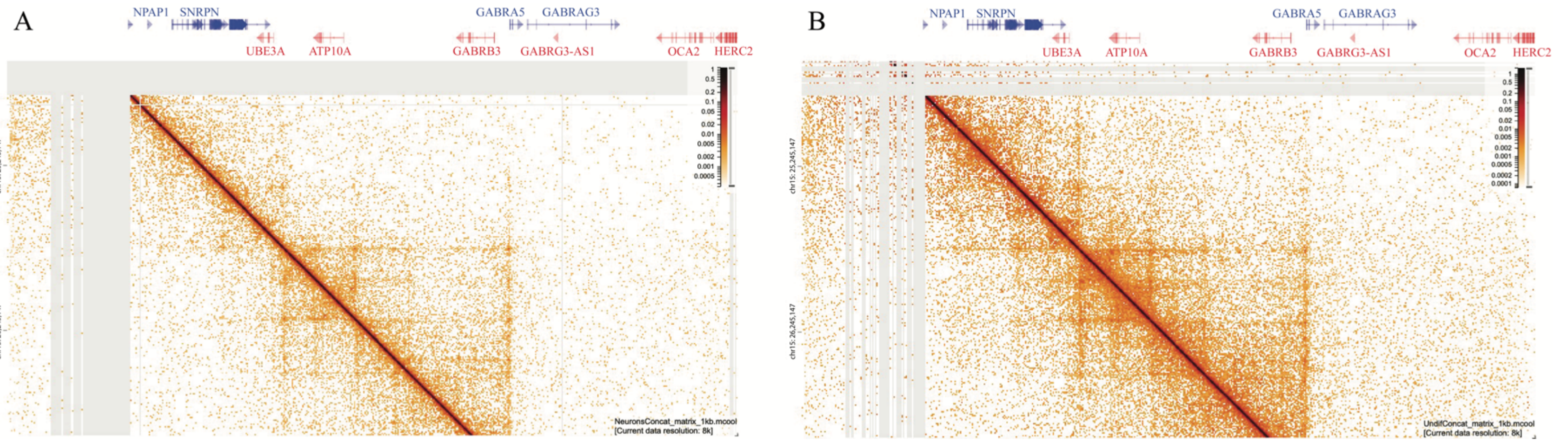


**Figure S2: HiChIP contact matrix.** HiChIP contact matrix generated using HiGlass(69), gray areas represent regions lacking contact (hg19; chr15:24073444-28488587 vs chr15:24611987-26881716) **A.** Neurons **B.** Undifferentiated LUHMES

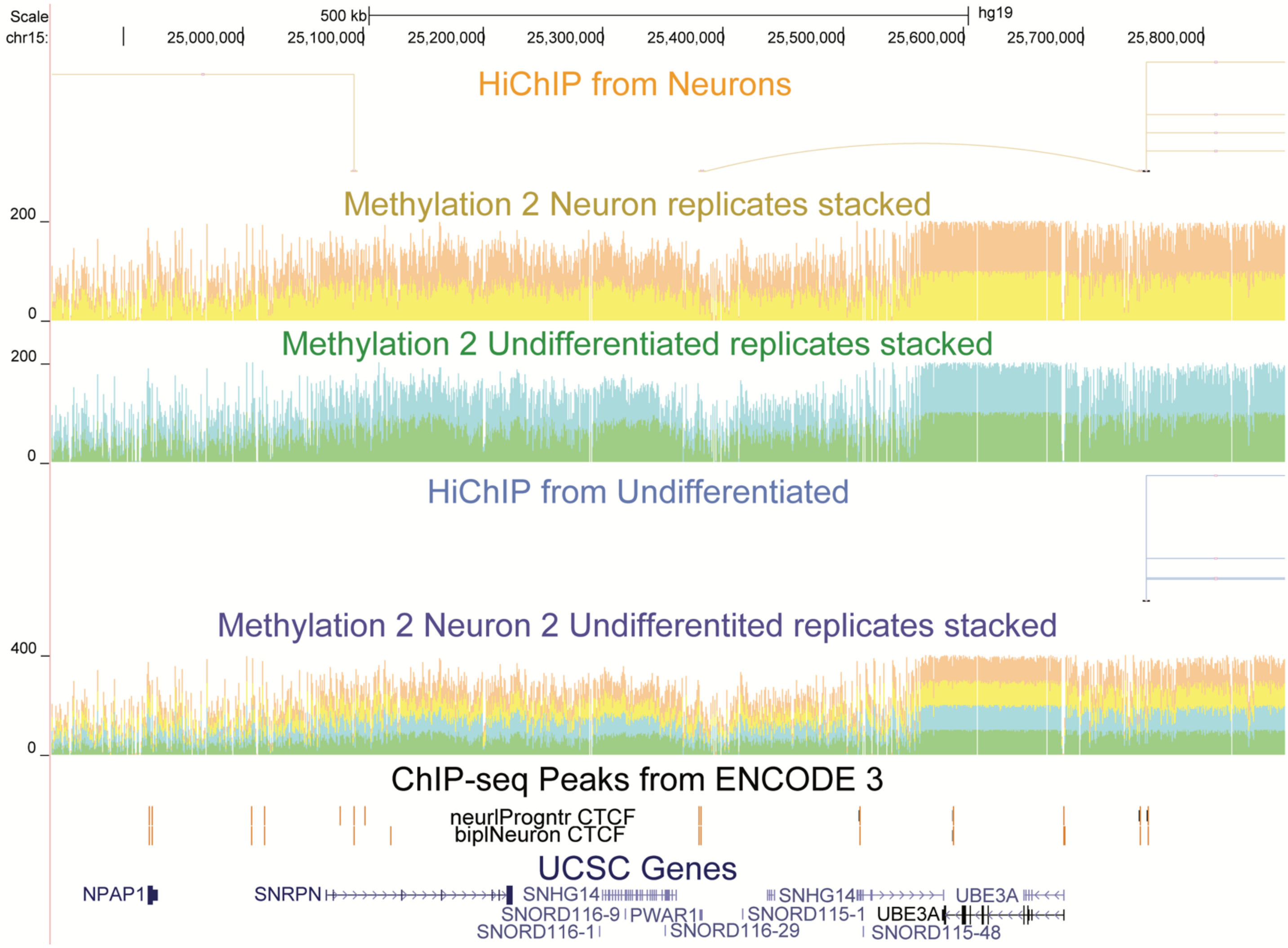


**Figure S3: LUHMES HiChIP and non-phased methylation profile.** Methylation from the positive strand, 2 replicates stacked and shown in different colors for contrast, values representing the sum of their percent methylation (max 200). A combined track stacking 2 neurons and 2 undifferentiated replicates (max 400). ChIP-seq Peaks from ENCODE 3 for CTCF in bipolar neuron and neuronal progenitor (hg19; chr15:23,832,378-25,962,021).

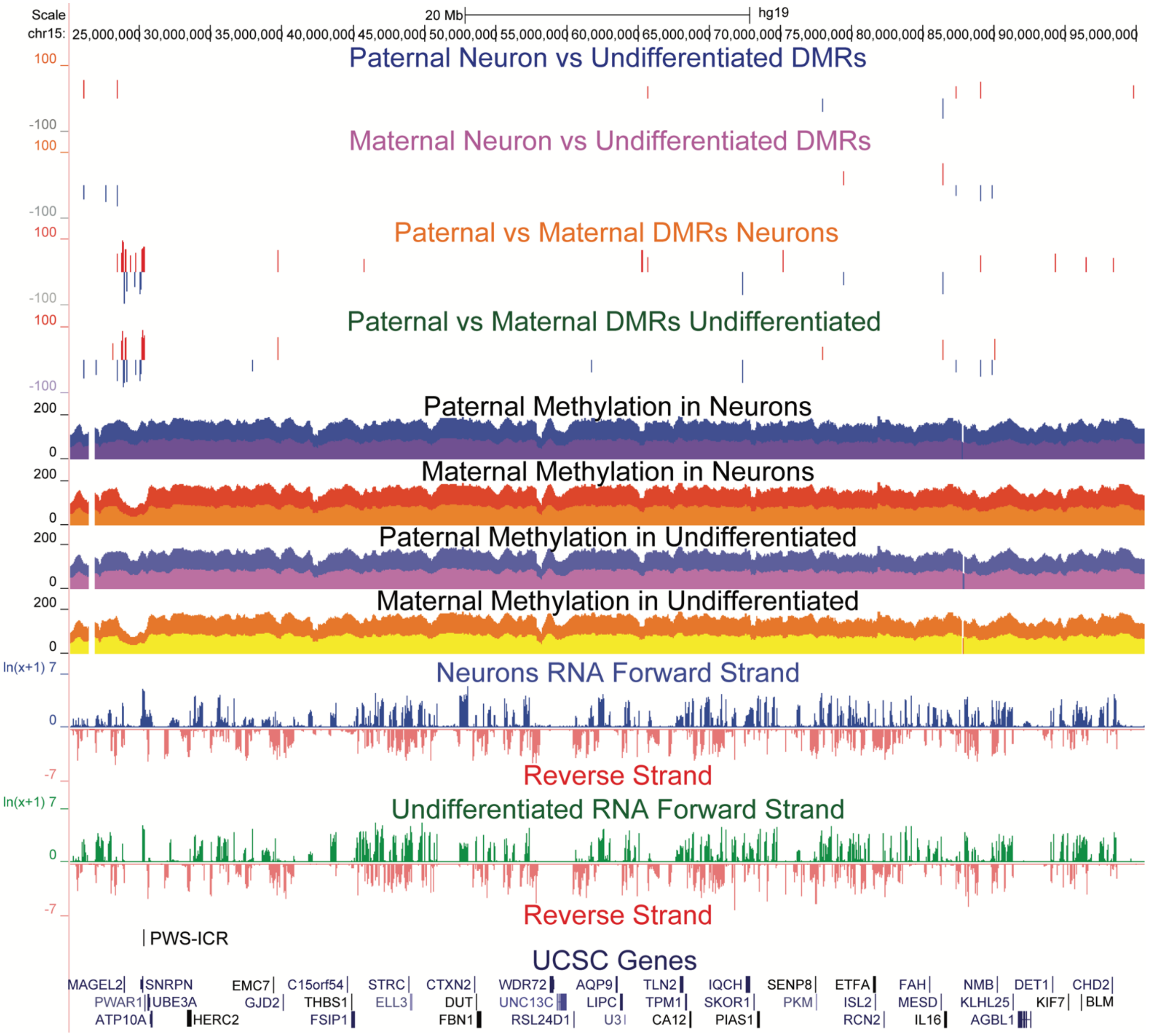


**Figure S4: Paternal and maternal DMRs for neurons vs undifferentiated LUHMES.** Few DMRs are seen when viewing at a large portion of chr15 comparing the paternal allele in neurons vs the paternal allele in undifferentiated cells as well as comparing the maternal allele in neurons vs the maternal allele in undifferentiated cells (hg19; chr15:20,141,900-95,560,739). We can also view how clustered the paternal vs maternal DMRs are for both cell types in the 15q11-q13 region.
